# Leucine-rich glioma inactivated 1 is a ganglioside-binding protein

**DOI:** 10.64898/2026.02.05.703988

**Authors:** Kévin Debreux, Christian Lévêque, Fodil Azzaz, Marion Sangiardi, Sarosh R Irani, Michael Seagar, Jacques Fantini, Oussama El Far

## Abstract

In LGI1-linked animal models of inherited autosomal dominant lateral temporal lobe epilepsy, increased neuronal excitability is accompanied by modifications in the AMPA/NMDA receptor ratio and a large decrease in Kv_1_ type potassium channels. However, the mechanism which links the absence of LGI1 to reduced expression of key neuronal ion channels is unknown. We observed multiple conserved canonical ganglioside-binding domains (GBDs) within human LGI1, mainly located in the EPTP domain. We show that GT1b is co-captured from native rat brain extracts by human LGI1 antibodies and, using SPR analysis, that recombinant full length LGI1 interacted with liposomes containing GT1b and GM1, but not GM3, lyso-lactosylceramide, phosphatidylserine or phosphatidylcholine. The ganglioside binding capacity of GBD peptide sequences exposed at the surface of LGI1 were confirmed using SPR and Langmuir film balance. Our data suggest that LGI1 interacts with gangliosides and may be involved in organizing lipid membrane platforms accommodating functional protein complexes. The loss of LGI1 could destabilize these platforms and contribute to reduced expression of key ion channels in *Lgi1*^-/-^ mice.

## INTRODUCTION

Leucine-rich glioma inactivated 1 (LGI1) is a secreted glycoprotein expressed in the central nervous system composed of four leucine rich repeats (LRR domain) and a β-propeller structure (EPTP domain), both mediating protein-protein interactions. The EPTP domain interacts with the ectodomain of membrane-inserted and catalytically-inactive members of the ADAM family of metalloproteases ADAM 22, 23 and 11 ^1 2 3 4^, and is a ligand for Nogo receptor 1. Direct interaction of LGI1 with ADAM 22 and 23 may be required for proper LGI1 trafficking and transport ^5^. Point mutations in LGI1, linked to inherited autosomal dominant lateral temporal lobe epilepsy [ADLTE] ^6 7^, result in neuronal hyperexcitability and epileptic seizures ^8 9 10^. These point mutations mostly inhibit LGI1 secretion ^11 12 13^ although some secreted mutants are impaired in their capacity to bind to ADAMs ^14 15^, suggesting that the phenotype of LGI1 secretion-defective or knockout animal models is linked to disruption of the ADAM22/LGI1 interaction ^16^. LGI1 deletion is associated with a decrease in both AMPA receptor-mediated synaptic transmission and AMPA / NMDA receptor ratio ^17 18 19^ as well as a large decrease in the expression of Kv1 channels ^20^. Neuronal hyperexcitability and the occurrence of lethal epileptic seizures ^8 9 10^ in *Lgi1*^*-/-*^ mice have been suggested to be primarily due to a major downregulation of Kv1 channels ^20^. Kv1 channels are present in complexes containing ADAM proteins and AMPA receptors, as well as MAGUKs and it has been hypothesized that LGI1 dimerization might form trans-synaptic complexes ^17 21 16^. The link between LGI1 and Kv1 channels is further highlighted in LGI1-antibody encephalitis ^22^, as patient autoantibodies against LGI1 co-immunoprecipitate Kv1 channels^22 23^.

The molecular mechanisms underlying the decrease in Kv1 expression have not yet been elucidated. LGI1 has been shown to modulate PSD95 incorporation at excitatory synapses ^17 19 21^. In heterologous systems, ADAM 22 and 23 induce opposite effects on Kv1 expression with ADAM 23 decreasing Kv1 currents independently of PSD95 and LGI1 while ADAM 22 potentiated Kv1 currents only in the presence of LGI1 ^24^. LGI1 interacts with ADAM 22 and 23, PSD95 putatively binds both Kv1 and ADAM 22 and the LGI1/ADAM interaction is likely important in PSD-95 function ^19^, hence the absence of LGI1 might reduce ADAM 22 / PSD95 clustering and contribute to destabilizing Kv1 at the membrane. However, ADAM-independent LGI1 interaction with specific membrane domains cannot be excluded especially since ADAM23 does not contain a PDZ-interacting domain^4^ and deletion of the PDZ-interacting domain of ADAM22 only partially reduces presynaptic Kv1 expression ^21^. Furthermore, the mechanism and function of juxtaparanodal Kv1 clustering is independent of PSD95 and ADAM, but depends on the presence of Caspr2 and TAG-1, a GPI-anchored protein associated with rafts. In neurons, LGI1 is secreted in both somatodendritic and axonal compartments and co-immunoprecipitates with juxtaparanodal proteins, including contactin1, TAG1, Caspr2 and Kv1 channel subunits ^25^. It is enriched at synaptic contact sites and is associated with oligodendrocytic as well as neuro-oligodendrocytic and astro-microglial protein complexes ^25^.

Gangliosides are sialylated glycosphingolipids enriched in the plasma membranes of the vertebrate nervous system. They are concentrated in membrane rafts, tightly packed with cholesterol and sphingomyelin, where they modulate the activity of cell surface proteins and constitute the binding sites for certain toxins and autoantibodies. Notably at the node of Ranvier, gangliosides participate in axo-glial contacts, including paranodal loop attachment to the axolemma, and contribute to the stability of potassium channel clusters in juxtaparanodes ^26^.

In the present report, prompted by the observation that LGI1 harbors several ganglioside-binding motifs, we asked whether LGI1 directly interacts with glycolipids, potentially providing an additional/alternative mechanism to modulate the expression and function of cell surface proteins.

## MATERIALS AND METHODS

### Reagents

1,2-Dimyristoyl-sn-glycero-3-phosphorylcholine (DMPC), monosialoganglioside (GM1), monosialoganglioside (GM3), disialoganglioside (GD1a), phosphatidylserine (PS), lyso-lactosylceramide were from Matreya LLC, GT1b (trisialoganglioside) was from Enzo Life Sciences. Anti LGI1 antibodies were previously described ^25^ and monoclonal anti GT1b was from Millipore.

### Expression plasmids

Construction of the inducible LGI1-expression plasmid has been previously described ^25^.

### Cell culture and LGI1 secretion

HEK293T cells were cultured at 37°C in DMEM (GlutaMAX) supplemented with 10% foetal bovine serum, 1% non-essential amino acids and 1% streptomycin/penicillin mix.

Transfection with LGI1-expressing plasmids was performed in optiMEM using PEI and LGI1 expression was induced by adding 300 ng/ml doxycycline 12 h after transfection. Secreted LGI1 was collected from the cell supernatant 48 h after induction. LGI1-containing optiMEM supernatant was cleared by centrifugation for 20 min at 16000xg and LGI1 was concentrated by ultrafiltration at 10000xg using Vivaspin 6 (30000 MWCO) tubes. Concentrated LGI1 was snap-frozen in liquid N_2_ and stored at -70°C.

### Peptides

Peptides from the rat LGI1 sequence (NP_665712.1) are listed in Table 1 and were synthetized by Genecust (France).

**Table 1.**
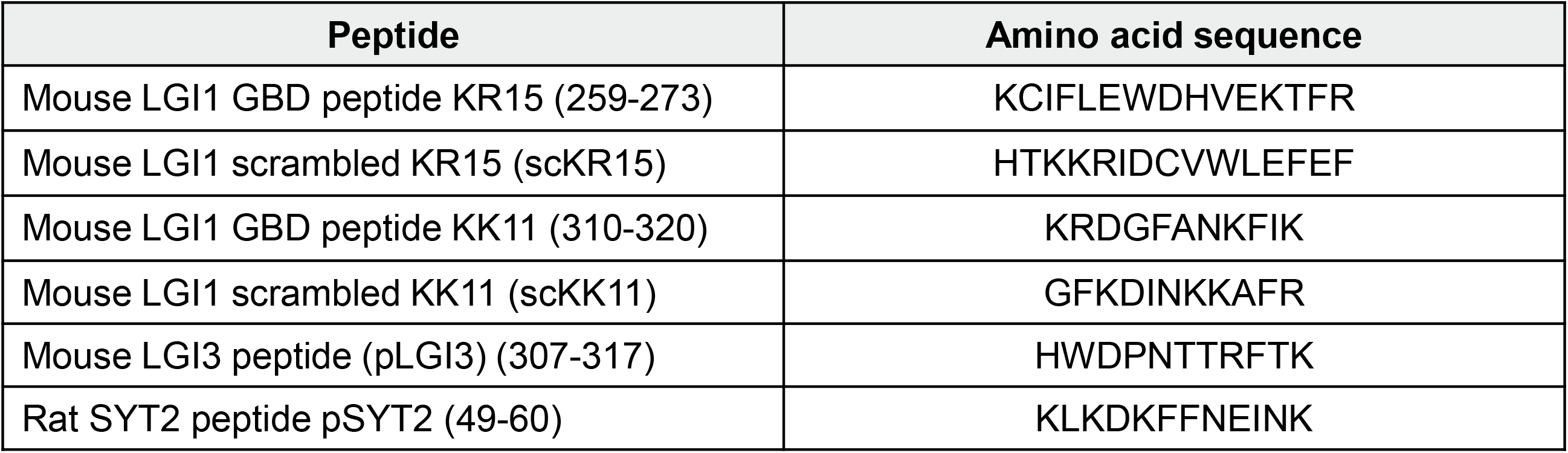
Amino acid sequences of peptides used in the study.

### Liposomes

Lipids were resuspended in chloroform / methanol (2vol/1vol) except for GT1b that was resuspended in methanol and stored at -20°C under nitrogen. Liposomes were prepared using mixtures of DMPC (1,2-dimyristoyl-sn-glycero-3-phosphocholine) with 8% GT1b, 8% GM1, 8% GM3, 8% lyso-lactosylceramide, 24% PS or 100% of DMPC for control liposomes. Solvent was evaporated under N2 and dried lipids (850 nmol) were resuspended by sonication in 1 ml of 25 mM Tris pH7.4, 150 mM NaCl. Liposomes were generated by extrusion at 55°C through 100 nm polycarbonate filters using a basic Lipofast apparatus (Avestin).

### Surface Plasmon Resonance

Recombinant anti-LGI1 IgG LRR3 ^25^ was covalently coupled at 25 °C on a Biacore T200 sensor chip CM5 (Cytiva) (circa 5000 RU) with an irrelevant human IgG on the control flow cell. 300 RU of LGI1 secreted by HEK293 cells and diluted in 10 mM HEPES/NaOH pH 7.4; 140 mM NaCl were then captured at a flow rate of 1 µl / min. Data were analysed using the Bievaluation2.0 software. To evaluate the specificity of lipid binding, liposomes of different compositions ^27^ were injected for 300 sec (10µL/min) over LGI1 captured on LRR3 and the control flow cell. Injections were separated by a regeneration step (6 sec HBS / 1% CHAPS wash followed by 2 x 6 sec injection of HBS both at 20 μl/min) and the injection of 100% DMPC liposomes. For experiments investigating LGI1 peptide binding to GM1, circa 7000 RU of liposomes containing 7% GM1 were immobilized on a L1 chip (Cytiva) whereas the control flow cell was functionalized with 100% DMPC at the same RU level.

### Co-capture of LGI1 and GT1b from native tissue

Rat brain synaptosomal fraction (P2) ^28^ was lysed to obtain a plasma membrane-enriched fraction (LP1) ^29^. LP1 was solubilized at 5 mg/ml in 25 mM Tris-Cl pH 7.4, 150 mM NaCl, 1% CHAPS, 5 mM EDTA and centrifuged at 100 000 g. A 3-fold dilution of the supernatant was injected over a control human IgGs or anti-LGI1 monoclonal antibody (LRR1^25^) covalently coupled to an SPR sensorchip before detection of binding partners by injection of specific antibodies.

### Langmuir balance

Peptide binding to gangliosides was determined using the Langmuir-film balance technique on a fully automated microtensiometer (μTrough SX; Kibron Inc.). Briefly, gangliosides resuspended in chloroform/methanol (1:1 vol/vol) were spread on a pure water subphase (800 μl) and interactions were monitored at 25 ± 1 °C by surface pressure modification of the monomolecular films. The initial surface pressure of ganglioside monolayers was set between 12 and 15 mN/m with an accuracy of measurement of ± 0.25 mN/m. Peptides were injected (final concentration 10 μM) into the aqueous phase underneath the ganglioside monolayer until equilibrium was reached with real-time measurements of changes in the kinetics of surface pressure. Data were collected and analyzed using the FilmWareX program (version 3.57; Kibron Inc.).

### Molecular Modeling

The full length LGI1 dimer was extracted from the crystal-coordinates in PDB file 5Y31 and the coordinates of the N-terminal part of LGI1 were generated by the online *Ab Initio* software Robetta (https://robetta.bakerlab.org/). The solvent-accessible surface of the LGI1 dimer was docked on the surface of a preassembled lipid raft system composed of 36 GT1b and 36 cholesterol molecules using “Visual Molecular Dynamics” (VMD) software ^30^. The initial coordinates and the PSF files of the cholesterol and gangliosides were obtained from the CHARMM-GUI website using the glycolipid and the bilayer modeler tools ^31^. The PSF file of the full length LGI1 dimer was generated using the Automatic PSF Builder tool on VMD. The LGI1/raft complex was minimized during 3000 steps using the molecular dynamics software NAMD version 2.14 for Windows 10 ^32^ coupled with the force field CHARMM36m designed for sphingolipids and carbohydrates ^33^. The images of the minimized complex were captured using PyMOL Molecular Graphics System ^34^. For the overview of the LGI1/raft complex, we used a raft inserted in a plasma membrane composed of 50% POPC and 50% cholesterol for the outer leaflet and 50% cholesterol, 30% POPS 20% POPE for the inner leaflet. This composition was chosen to mimic a neuronal membrane.

## RESULTS

### Identification of ganglioside binding motifs in LGI1

Studies of protein/lipid interactions identified short amino acid sequences in beta-amyloid and alpha-synuclein which then yielded a consensus motif [K/H/R-x(1-4)-Y/F-x(4-5)-K/H/R] to predict putative GBDs (ganglioside binding domains) in proteins ^35^. Analysis of the human LGI1 sequence revealed multiple and highly conserved (human to mice) canonical GBDs mainly located in the EPTP domain (Figure 1A-D), often at the junction between two β-propeller blades (Figure 1A), and 1 in the hinge region linking the LRR and EPTP domains. These sequences are poorly conserved in other LGI family members (Supplementary Figure 1). Most exposed GBD are highlighted in Figure 1D and served to design the experimental peptides.

**Figure 1.**
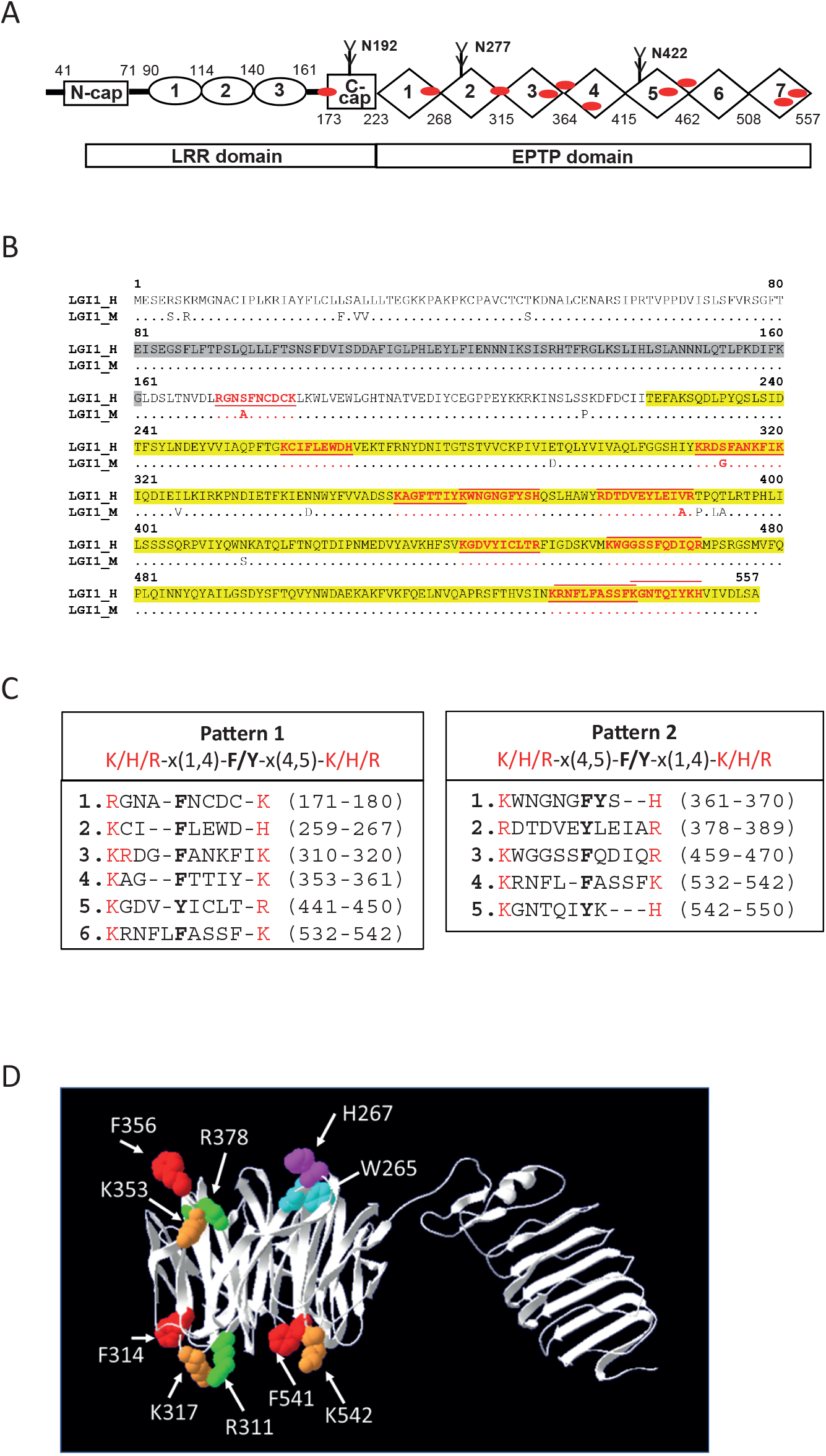
Consensus ganglioside-binding domains in LGI1. **A**. Schematic structure of LGI1. LGI1 domains are shown as ovals (LRR) and lozenges (EPTP), with N-glycosylation sites (branched lines) and approximate positions of putative GBDs (red marks). **B**. Sequence alignment of human and mouse LGI1. GBD sequences are in bold red and underlined (Pattern 1 in C) or overlined (Pattern 2 in C). LRR and EPTP domain sequences are highlighted in grey and yellow respectively. **C**. GBDs in mouse LGI1. Both N toward C-terminal (Pattern 1) (K/H/R-x(1-4)-Y/F-x(4-5)-K/H/R) and C towards N-terminal (Pattern 2) (K/H/R-x(4-5)-Y/F-x(1-4)-K/H/R) GBDs were identified using the online Prabi-Gerland tool (https://npsa-prabi.ibcp.fr/cgi-bin/npsa_automat.pl?page=/NPSA/npsa_pattinprot.html). **D**. Position of certain principal amino acids of GBDs exposed at the surface of the EPTP domain of human LGI1.

### Co-capture of LGI1 and GT1b from native tissue

To verify if gangliosides are present in native LGI1 membrane complexes, solubilized rat brain plasma membrane fractions (LP1) were injected over an anti-LGI1 monoclonal antibody (LRR1 ^25^) previously immobilized on an SPR sensor chip. LGI1 was specifically and stably captured in addition to ADAM22 and 23, its two well characterized partners (Supplementary Figure 2). An anti-GT1b antibody verified GT1b presence in the LGI1-linked molecular complex.

### LGI1 interaction with complex gangliosides

To test whether full length LGI1 can bind gangliosides, recombinant LGI1 was recovered from the culture medium of transfected HEK293 cells and then immobilized by specific capture on an LGI1 (LRR3) monoclonal antibody ^36 25^ covalently coupled to an SPR sensor chip. DMPC liposomes containing different gangliosides or phosphatidylserine (PS) were then injected into the flow cell and showed a specific interaction between LGI1 and liposomes containing GM1 or GT1b, but not liposomes containing GM3, lysolactosylceramide or PS or pure DMPC (Figure 2A-B). This result indicated LGI1 interacts with complex gangliosides harboring a tetraose backbone.

**Figure 2.**
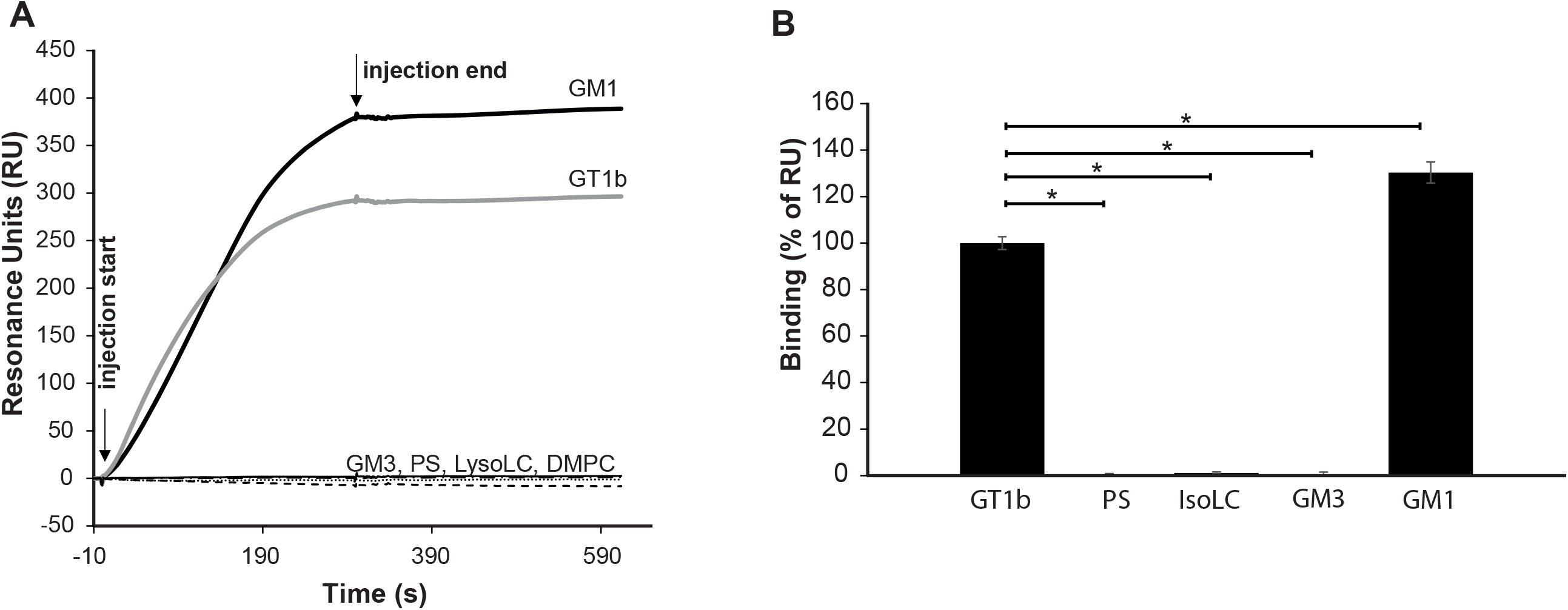
SPR analysis of the interaction between LGI1 and GM1 or GT1b gangliosides. A. Sensorgrams illustrating the binding of DMPC liposomes containing GM1 (black trace) or GT1b (grey trace) to recombinant LGI1 immobilized on the chip surface via a coupled human LGI1 monoclonal antibody. Non-specific binding to a parallel flow cell carrying irrelevant human IgG has been subtracted. DMPC liposomes (- - -) containing GM3 (…), lysolactosylceramide (IsoLC; thin black trace), or DOPS (-.-) did not display any specific binding. **B**. Histograms representing the specific binding of gangliosides (GT1b n=13; PS n=7; IsoLC n=7; GM3 n=7; GM1 n=3) to pre-immobilized LGI1 as illustrated in A. Background binding corresponding to binding of 100% DMPC has been subtracted. Results were normalized to data obtained from injections of GT1b-containing DMPC liposomes. Statistical analysis was performed using Wilcoxon Mann & Whitney. n represents the number of determinations cumulated from at least three independent experiments.

### Interaction of LGI1-derived peptides with gangliosides

To confirm that gangliosides directly interact with LGI1, we assessed ganglioside binding of two distinct GBD containing sequences in LGI1 (Figure 1 C, D; Table 1). Among the consensus GBD, we chose two surface-exposed motifs that are located in the EAR sub-domain of the EPTP region, outside of the LGI1/ADAM22 interface ^1^. We synthetized the corresponding peptide sequences KR15 (259-273) and KK11 (310-321) as well as their cognate scrambled versions (scKR15 and scKK11; Table 1) and assessed their binding capacity to GM1-containing liposomes with SPR. KR15 and KK11 interacted with immobilized GM1-containing liposomes (Figure 3) similar to observations we have reported with the ganglioside-binding pSYT2 peptide ^27^. Scrambling these peptides significantly decreased these interactions and the peptide corresponding to KK11 in LGI3 (pLGI3, Table 1), which is devoid of a GBD consensus, did not show any binding (Figure 3A). To investigate the interactions of these peptides with gangliosides, we used a Langmuir monolayer approach which showed that introducing KK11 into the aqueous phase resulted in a significant increase in initial surface pressure with GM1 monolayers whereas the signal was significantly delayed using scKK11 (Figure 3 B). KR15’s interaction with GM1 could not be measured using this method due to its spontaneous surface activity. To measure the specificity of the interaction, monolayer films of GM1 and GT1b were prepared at varied initial pressures and the maximal increase in surface pressure was monitored at equilibrium plateau. The increase in surface pressure induced by peptide KK11 diminished proportionally with the increase in ganglioside density at the water / air interface because the insertion of the peptide within the lipid monolayer is easier (low resistance) at low surface pressure (Figure 3C). The linearized variations in pressure displayed a gentler slope with GM1 than GT1b, with critical pressures of insertion extrapolated at the x-axis intercept at 42 mM/m and 29.3 mN/m, for GM1 and GT1b respectively. These values represent the critical pressure of insertion in the monolayers. The value for GT1b is close to the lipid density in cell membrane (30 mN/m) whereas the higher value for GM1 indicated a stronger affinity. In contrast to KK11, scKK11 did not show an exploitable binding pattern (Figure 3D) due to point dispersion reflecting low binding affinity. These results show that both LGI1 peptides interact individually with complex gangliosides with a higher affinity for GM1 than for GT1b and suggest that their corresponding domains are involved in LGI1 binding to membranes.

**Figure 3.**
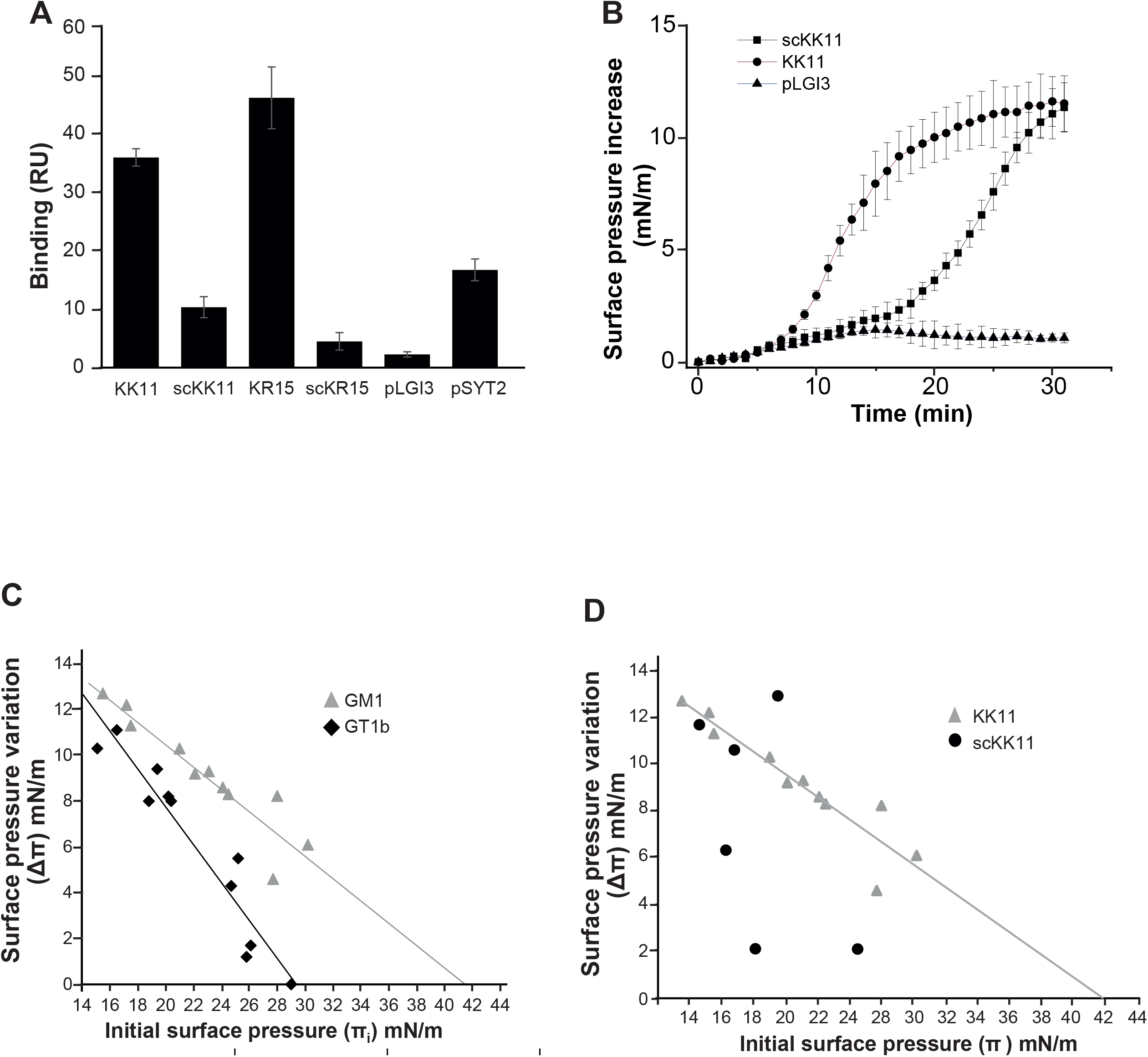
Interaction of LGI1 peptides with gangliosides. **A**. SPR measurement of the LGI1 peptides KK11 and KR15 binding to GM1-containing liposomes. Peptides were injected at 30 µM for 3 minutes over a sensorchip functionalized with liposomes containing 93 % DMPC and 7 % GM1 in the experimental flow cell and 100 % DMPC in the control flow cell. Scrambled LGI1 peptides (scKK11 and scKR15), an LGI3 peptide corresponding to the position of KK11 as well as a ganglioside-binding synaptotagmin 2 peptide were used as controls. Non-specific binding on the control flow cell has been subtracted and specific signals at 5s after the beginning of the injection are shown. Results are ± SD from three independent experiments. **B**. Kinetics of KK11, scKK11 and pLGI3 interaction with a GM1 monolayer. Represented measurements of surface pressure increase are mean of triplicates ± SD **C**. Interactions of peptides KK11 (grey triangles) or scrambled scKK11 (black circles) with a GM1 monolayer were analyzed as in (**A**). Linearization of the data yielded a linear fit with KK11 (y = -0.48x + 20.05) with R^2^ = 0.86. Measurement points with scKK11 were dispersed and the linear fit is not represented since it yielded an R^2^ = 0.27 **D**. Interactions of peptides KK11 with GM1 (grey triangles, y = -0.48x + 20.05; R^2^ = 0.86) or GT1b (black diamonds, y = -0.83x + 24.3; R^2^ = 0.91) monolayers.

### Molecular dynamics of LGI1 interaction with a GT1b raft

In order to explore the theoretical structural correlates of LGI1 binding to gangliosides via exposed LGI1 GBDs, we carried out molecular modeling and assessed the propensity of KK11 and KR15 peptides (Figure 1 C, D; Table 1) to bind to a GT1b raft in a membrane context.

The crystal structure of LGI1 was extracted from LGI1/ADAM22 complex PDB file 5Y31 ^1^. An LGI1 dimer was then docked at the surface of a GT1b raft inserted in a lipid bilayer and the structure of the complex was minimized (Figure 4). Since the LGI1 structure was extracted from a complex with ADAM22, we oriented the LGI1 ADAM22-free interface towards the GT1b raft. As shown in Figure 4, the molecular dynamics of this structure is compatible with an interaction between LGI1 and the GT1b raft via a zone corresponding to peptides KR15 (259-273) and KK11 (aa 310 – 320).

**Figure 4.**
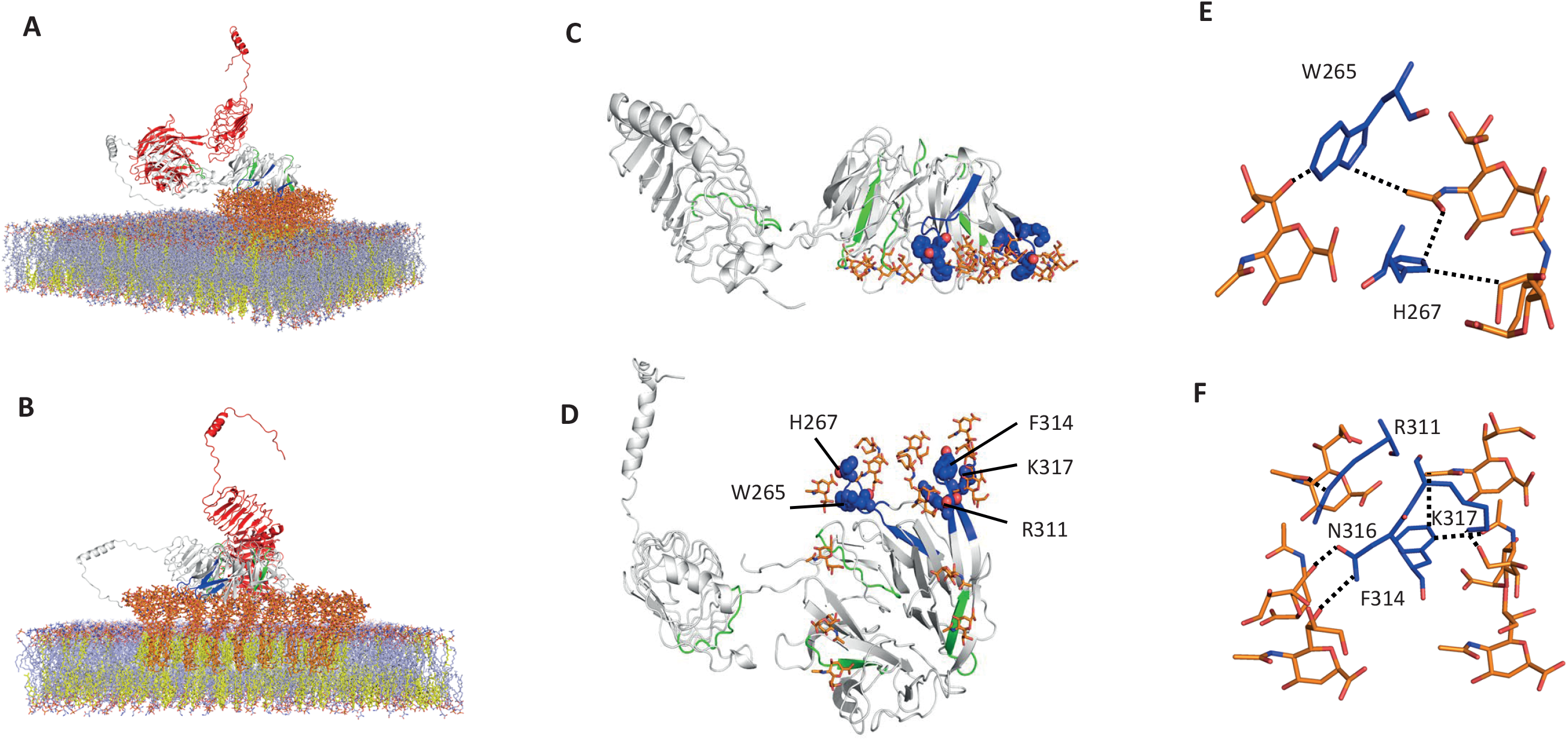
Molecular dynamics analysis of LGI1/gangliosides interactions. **A**. Overview of the LGI1/GT1b raft complex embedded in a plasma membrane composed of 50% POPC and 50% cholesterol for the outer leaflet and 50% cholesterol, 30% POPS 20% POPE for the inner leaflet, a composition chosen to mimic a neuronal membrane. The LGI1 dimer is shown as a cartoon (red and white for monomers 1 and 2 respectively), the lipids are shown as sticks (light blue for phospholipid and yellow for the cholesterol) with the glycolipids displayed as orange sticks. On monomer 2, the GBDs are shown in green and blue for the KK11 peptide that was studied experimentally. **B**. A section of the membrane to obtain an insight into the GBDs in contact with the lipid raft. **C, D**. A top and front snapshot (respectively) of the LGI1 dimer to highlight accessible surface in the lipid raft. **E and F**. Atomic details of the interaction between the sialic acids and key amino acids of the GBDs. The interaction strength is principally conferred by CH-pi and hydrogen bonds.

## DISCUSSION

Short amino acid sequences, initially discovered in beta-amyloid and alpha-synuclein, have led to identification of a consensus motif to predict protein GBDs^35^. The majority of the putative GBDs we detected in the LGI1 sequence were located within the EPTP domain which interacts with ADAM22 ^1^. When LGI1 is bound to ADAM22 its flexibility is too restricted, and it is positioned too far from the membrane bilayer to permit cis interactions with gangliosides to occur. However, LGI1 can form homodimers and homomultimers^37^, which could theoretically form cis or trans interactions with lipids.

The linear sequence comparison of LGI1 with the other LGI isoforms (LGI 2, 3 and 4) showed (Supplementary Figure 1) that the majority of the theoretical GBDs in LGI1 are not conserved. A direct investigation of ganglioside binding to other LGIs should however be performed in order to establish this finding.

The hypothesis that GBDs in LGI1 can bind to gangliosides was first evaluated experimentally, using SPR analysis. Recombinant full length LGI1 captured native GT1b from solubilized brain extracts and was shown to interact with DMPC liposomes containing GM1 or GT1b but not GM3, lyso-lactosylceramide or PS. PS did not support binding, even though added at a concentration providing equivalent electrostatic charge to GT1b. Furthermore, LGI1 bound to GM1 but not to GM3, although both contain a single sialic acid. Consequently, binding does not depend uniquely on simple electrostatic interactions, but also displays specific structural requirements involving the glycoside chain and the relative positions of the negative charges of sialic acid in relation to this chain. In the context of LGI1-associated pathologies, it is interesting to note that a defect in ganglioside synthesis has been shown to be associated with epilepsy ^38 39^. The complex ganglioside GM1 is enriched in synaptosomes ^40 41^ and has been suggested to play a role in determining axonal fate ^42^ and together with GT1b is implicated in glial / neuronal contact sites ^43^. Compared to GM1 and GT1b, the simple ganglioside GM3 that fails to interact with LGI1 is a prominent component of non-neuronal cells and is particularly enriched in oligodendrocytes. It was suggested to be involved in transmembrane signaling, cell adhesion and motility through interaction with integrins ^44^ and was earlier shown to be localized mostly at the somatic level ^43^. GT1b in axonal microdomains participates in axon-myelin stability by directly interacting with the NgR1 ligand MAG (myelin associated glycoprotein) ^45 46^. This is particularly interesting in view of the distribution of LGI1 and its associated molecular complexes ^25^ and that NgR1 directly interacts with LGI1^47^. These results suggest that LGI1 binding to gangliosides may be important in organizing certain molecular complexes restricted to specific neuronal microdomains.

SPR experiments confirmed the ganglioside-binding capacity of two surface exposed GBD sequences in the EPTP region of LGI1. This binding was sequence-dependent since peptides with scrambled sequences showed a drastically decreased binding capacity. Also, a peptide sequence corresponding to the position of KK11 in LGI3 did not show GBD sequence conservation and ganglioside binding. The Langmuir monolayer approach was used to evaluate further the ganglioside-binding properties of the KK11 peptide and confirmed interactions with both GM1 and GT1b. As suggested by the SPR experiments, scKK11 binding to gangliosides was much perturbed and the determination of its critical insertion pressure in the GM1 monolayer was not possible. The absence of interaction of the LGI3 peptide with gangliosides does not exclude a global general feature of LGI proteins for interaction with gangliosides. Indeed, several GBD domains are present in LGI3 sequence. In addition, it was previously reported that Aβ upregulates the expression of LGI3 and that the latter colocalizes with endocytosis-associated proteins and lipid rafts markers on neurons and astrocytes ^48^ suggesting that LGI3 interacts with raft components.

Molecular modeling confirmed that in the context of a full-length 3D structure of an LGI1 dimer, the two LGI1 GBD sequences experimentally tested in this study can interact with clusters of GT1b molecules in a membrane raft environment.

Taken together our data are consistent with the association of LGI1 with ganglioside-rich membranes. ADAM22, the protein receptor for LGI1^1^ also interacts with TAG-1 ^49^, a protein which promotes membrane expression and clustering of GM1 ^50^. These properties may favor the assembly of lipid rafts rich in GM1, associated with ADAM22 and LGI1. LGI1 tetramers ^1^ and hexamers ^37^ could contribute to the formation of a ganglioside-rich environment, which stabilizes membrane protein complexes, including Kv1 channels. Our recent finding of Kv_2.1_ downregulation in *Lgi1*^*-/-*51^ argue also in this direction since Kv_2.1_ was shown to bind to Flotilin ^52^, a raft associated protein.

Modifications in membrane tension influence membrane dynamics and may impact the stability of certain proteins ^53^. Actin cytoskeleton polymerization and reorganization are modulated by Rho GTPase ^54^ activity of which was shown to be modified in *Lgi1*^*-/-*^ . LGI1 may therefore keep a certain tension on the membrane and upon LGI1 deletion, changes in membrane tension may modify Rho signalling et actin cytoskeleton reorganization influencing endocytosis from certain membrane domains.

Further studies will be necessary to address the biological relevance of ganglioside-binding by LGI1. These will involve determining whether interactions of LGI1 with the plasma membrane of cultured cells, in the absence of its canonical receptors ADAM22, 23 or 11, is inhibited by ganglioside depletion and subsequently rescued by addition of extrinsic gangliosides. Identification of the GBDs involved will then require directed mutation in full length LGI1. However, this approach may be hindered by the fact that many point mutations in LGI1 inhibit its secretion.

In conclusion our findings point to a novel mechanism by which LGI1 homo-oligomers could interact with rafts in specific membrane sub-domains of neurons and glial cells, although the functional significance has yet to be explored.

## Funding

This work was supported by the Institut National de la Santé et de la Recherche Médicale (INSERM), Aix-Marseille Université (AMU) and the Agence Nationale de la Recherche (ANR) (grant ANR-17-CE16-0022). The PhD thesis of K.D. was supported by a fellowship from the French Ministry of Research (MESRI).

---

**Supplemental Figure 1.**
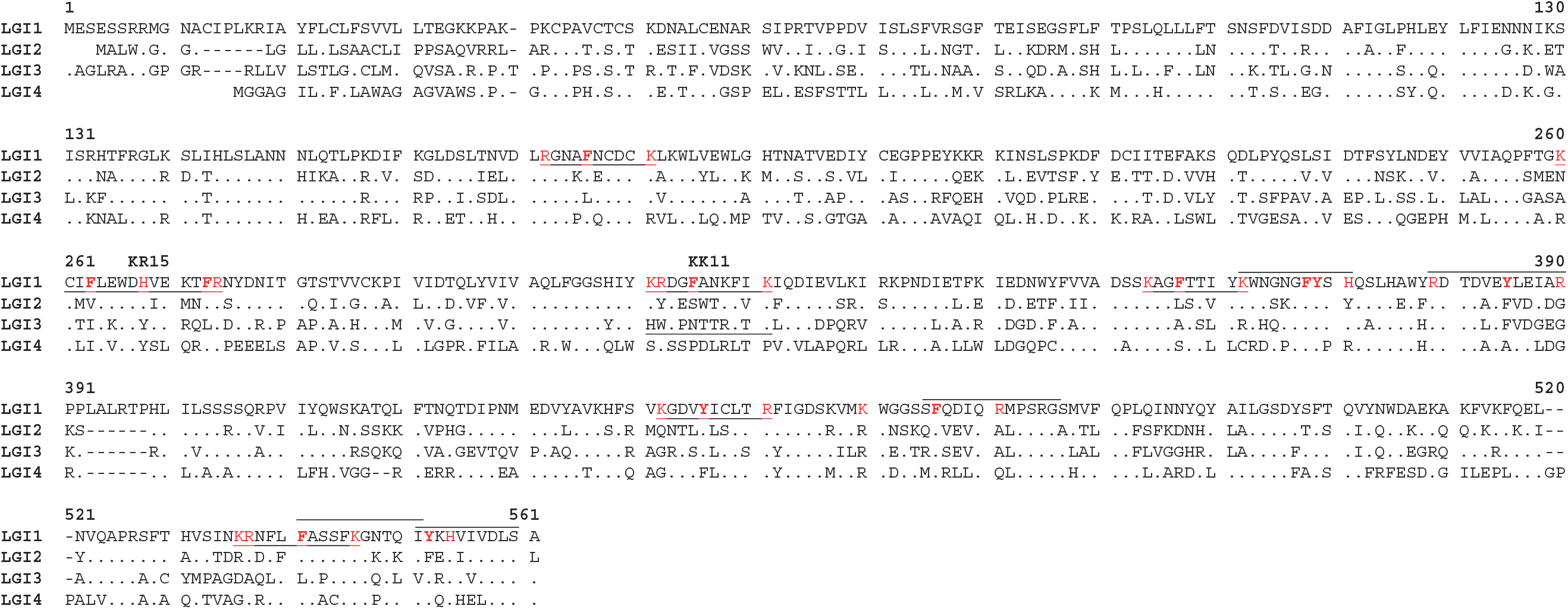
Alignment of the amino acid sequences of mouse LGI1, 2, 3 and 4. Conserved residues are marked by dots. GBDs described in Figure 1 are underlined (Pattern 1) and overlined (Pattern 2). Positively charged amino acids delimiting the GBDs are in red and central hydrophobic residues of GBDs are in red and bold. KK11 and KR15 indicate the position of the GBD binding peptides used in this study. The corresponding LGI3 peptide (pLGI3) is underlined

**Supplemental Figure 2.**
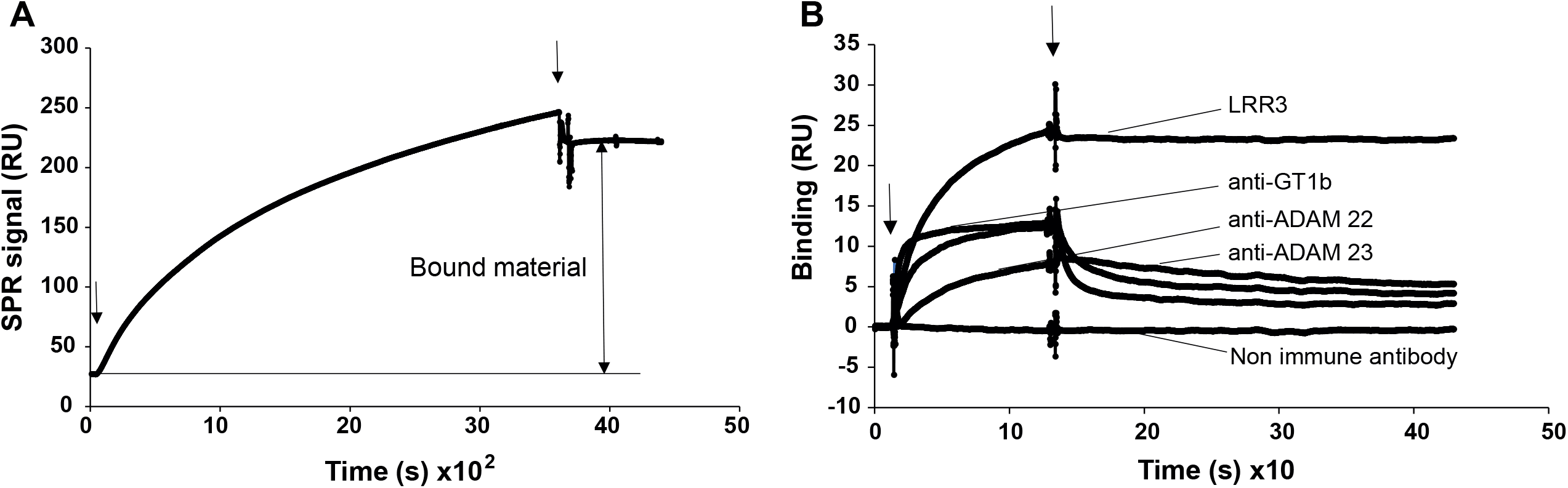
Detection of GT1b associated with LGI1 from LP1 extract. **A**. LP1 was solubilized in CHAPS and injected over anti-LGI1 and control antibody **B**. Antibodies directed against GT1b, LGI1 (LRR1 and LRR3) and ADAM 22 or 23 (10 µg/ml) were injected for 2 minutes at 10 µl/min. Arrows indicate start and end of injections. Non-specific signals on non-immune human IgGs have been subtracted from the presented curves.

**Table 1**

List of peptides used from the rat LGI1 sequence (NP_665712.1)

## Notes

### Competing Interest Statement

The authors have declared no competing interest.

